# T cell-Macrophage Interactions Potentially Influence Chemotherapeutic Response in Ovarian Cancer Patients

**DOI:** 10.64898/2026.03.02.709041

**Authors:** Sodiq A. Hameed, Walter Kolch, Vadim Zhernovkov

## Abstract

Tumor development and progression involve complex cell-cell interactions and dynamic co-evolution between cancer cells, immune cells and stromal cells in the tumour microenvironment, and this may influence therapeutic resistance. A large proportion of this network relies on direct physical interactions between cells, particularly T-cell mediated interactions. Cell-cell communication inference has now become routine in downstream scRNAseq analysis, but this mostly fails to capture physical cell-cell interactions due to tissue dissociation. Doublets occur naturally in scRNA-seq and are usually excluded from analysis. However, they may represent directly interacting cells that remain undissociated during library preparation. In the present study, we uncover the physical interaction landscape of the ovarian tumour microenvironment using the scRNAseq datasets from 13 treatment-naive ovarian cancer patients. Focusing on T-cell-Macrophage (T-Mac) interaction doublet, we reveal the modulatory effect of macrophages on T cells and the potential influence of this interaction on therapeutic response. Our findings show that T-Macs from resistant patients are functionally polarized to the M2 phenotype and engage T cells to induce T-cell exhaustion. Whereas, T-Macs from sensitive patients are predominantly of the M1 polarized phenotype, physically engaging T cells that lack exhaustion signatures. We also demonstrate that T cells and macrophages in T-Mac doublet are interacting primarily for the purpose of antigen presentation, with the enrichment of several ligand-receptor pairs involved in TCR-MHC interactions and immune synapse formations. We partly validated some of these findings from a spatial transcriptomics dataset of ovarian cancer patients from a separate cohort.

## 1.0 INTRODUCTION

Ovarian cancer remains the leading cause of gynaecological related cancer deaths in women globally, typically presenting with non-specific symptoms such as abdominal distension and abdominal pain [1]. It comprises a heterogeneous group of diseases that can be subdivided into at least 3 main subtypes, including clear cell, endometrioid, and high-grade serous ovarian cancer (HGSOC), each with distinct histological and molecular features [2]. HGSOC is the most common subtype that accounts for nearly 80% of all ovarian cancer related deaths [3]. Ovarian cancer is usually diagnosed at a late advanced stage due to the ambiguous symptoms and lack of definitive screening strategy [4], with only 20% of the disease being diagnosed at early stages for successful treatment [3]. HGSOC patients often present advanced stage diseases involving pelvic and peritoneal cavities with malignant ascites, and sometimes transcoelomic and distant metastases[1]. The standard treatment for newly diagnosed disease includes cytoreductive surgery and platinum-based chemotherapy [4]. Although many patients initially respond, recurrence is common and long-term survival remains poor, largely due to acquired chemoresistance and extensive inter- and intra-tumour heterogeneity. Thus, limiting the 5-year survival rate to around 30% with 75% of patients dying from the disease within 5 years of diagnosis . Increasing evidence suggests that cellular heterogeneity within the tumour microenvironment, including malignant, stromal, and immune compartments, contributes to therapeutic resistance, but the relevant cellular states and interactions remain incompletely understood [5].

Cell-cell communication is a fundamental phenomenon that underlies diverse tissue structure, function, and processes in homeostasis, including immune responses. These interactions are dependent on complex biochemical signalling pathways that regulate individual cell processes and intercellular interactions [6]. Cells in multicellular organisms continuously communicate from early embryonic development to adult life through these dynamic interaction networks. Tumor development and progression involve complex cell-cell interactions and dynamic co-evolution between cancer cells, immune cells and stromal cells in the tumour microenvironment and this may influence therapeutic resistance [7]. Cell-cell communications are mediated by a wide range of factors, some of which support structural interactions which bring about physical cell-cell interactions, while others mediate cell communications over short or long distances via secreted factors [8]. Single-cell RNA sequencing (scRNA-seq) enables high-resolution profiling of tumour ecosystems and has motivated computational inference of intercellular communication, typically via ligand-receptor expression patterns [9]. [10] [11] [12] [13]. However, conventional scRNA-seq requires tissue dissociation and therefore largely removes information about physical cell-cell contacts. Approaches that partially preserve tissue structure can recover contact information but are often technically demanding [14] [15] [16] [17] [18] [19]. Newer tools, from our group and others, such as Neighbor-seq [20], CIcADA [21], and more recently ULMnet [22], exploit naturally occurring biological doublets (or multiplets) in scRNAseq to infer physical cell-cell interactions from conventional scRNAseq datasets.

Physical cell-cell interactions are very crucial in cancers including ovarian cancer, particularly in immune responses involving T cells where they physically contact antigen-presenting cells (APCs), such as macrophages, loaded with tumour antigens. Following antigen presentation, primed T cells then go on to physically contact and kill cancer cells expressing the tumour antigens [23]. In the present study, we applied ULMnet [22] to infer physical cell-cell interactions in a scRNAseq dataset from 14 ovarian cancer patients. Focusing on antigen presentation dynamics, we characterize how T cells interact with macrophages in ovarian cancer patients and assess how these dynamic interaction patterns relate to therapeutic response.

## 2.0 METHODS

### 2.1 Single-cell RNA sequencing and spatial transcriptomics data analysis

A publicly available preprocessed ovarian cancer scRNAseq dataset and associated metadata, including cell annotations and clinical information, were obtained from a study by Zheng et al. [24]. In addition, raw count matrices for the same samples, with doublets retained, were obtained from the study authors. Concurrently, the unprocessed raw count matrices (doublet present) of the same dataset were privately obtained from the authors. This study consisted of 14 ovarian cancer patients, 13 of which were treatment naive, undergoing debulking surgery, and subsequently 6-month course of platinum-based adjuvant chemotherapy. Fresh samples were taken during debulking surgery to include primary ovarian tumor, omental metastasis, primary lymph node, malignant ascites and peripheral blood. Patients whose disease progressed within 6 months after the completion of adjuvant therapy were classified as “resistant” while the others were classified as “sensitive”. Notably, there were 4 resistant and 9 sensitive patients [24]. Datasets were processed using the Seurat package v5.1.1 in R v4.2.2. The raw scRNAseq count matrices were preprocessed to include only cells containing a minimum of 200 genes and less than 10 percent mitochondrial genes. Next, both scRNAseq datasets were log2 transformed, scaled, and the top 2,000 highly variable genes (HVG) were used for downstream analysis. A principal component analysis (PCA) was performed on the top 2,000 HVGs, and the top 40 principal components (PCs) were selected based on visualization from an elbow plot of the first 50 PCs to generate a uniform manifold approximation and projection (UMAP) plot. Then, cells were clustered using the nearest neighbor clustering algorithm at a resolution of 0.3. Batch effects were corrected, across patients and tumour types, using the Harmony batch correction algorithm [25].

A second publicly available preprocessed primary ovarian cancer scRNAseq count matrix and the associated metadata, including cell annotations, from a study by Desienko et al. [26] were obtained from the GEO database (GSE211956). This also included the spatial transcriptomics data of primary ovarian cancer tissue sections from 8 patients, containing spot-level count matrix, spatial imaging data and tissue coordinates. Samples were taken during interval debulking surgery with patients undergoing 3-5 cycles of neoadjuvant chemotherapy, resulting in 3 patients with good response, 2 patients with partial response, and 3 patients with poor response to chemotherapy, based on chemotherapy response scores [26]. For scRNAseq, the data was processed to include only genes present in at least 3 cells and only cells with a minimum of 500 genes, less than 15% mitochondrial genes, and more than 500 total UMI counts. Data was processed as above, but using 30 PCs for UMAP visualization and a resolution of 0.35 for clustering. Cells were assigned cell type labels by transferring the annotations present in the obtained metadata. For spatial transcriptomics, the individual matrices and imaging data from the 8 patients were loaded to create 8 spatial objects. Next, data were normalized and transformed using the SCT method [27] and PCA was performed on each spatial object after identifying the top 3000 HVGs. Spatial spots were then clustered using the nearest neighbor clustering approach, followed by UMAP projection and visualization of the first 30 PCs. To decipher the possible cellular composition of spatial spots, a spot-level deconvolution was performed using the robust cell type decomposition (RCTD) method from the Spacerx R package [28], utilizing the scRNAseq data from the same study as reference.

### 2.2 Doublet predictions: T-Macrophage doublet isolation

Doublet predictions and inferences were performed using the ULMnet R package [22]. Briefly, this method uses univariate linear regression models to score each cell in a scRNAseq data for signature expressions. Cells with single positive signature expressions would be labelled singlets while cells with dual or multiple positive signatures would be labelled doublets or multiplets. We applied ULMnet to the ovarian cancer datasets from Zheng et al. [24], that were processed in section 2.1. First, the annotated singlet dataset (doublet removed) was used for signature generation, using the GetSignature function with the default parameters (n= 100, p_val = 0.05). This generates a signature of 100 genes for each cell type in the annotated cell clusters by performing a one-vs-all differential expression (DE) analysis, ordering DE genes by logFC, and selecting the top 100 genes using a p value cutoff of 0.05. Next, the generated signatures from the annotated singlet scRNAseq data were then used to predict each cell in the unannotated scRNAseq data (doublet present) through a series of steps. ULMnet first models the cell type-specific signatures over the gene expression of each cell in the unannotated data to assign signature scores, using univariate linear models. Then each cell is assigned enrichment labels based on the signature scores using the GetCellAssignment function with the default parameters (p_val = 0.05, cut_off = 1). Cells with positive enrichment score for only one cell are labelled singlets while cells with more than one positive signature enrichment score are labelled multiplets (for example, doublets for dual signatures, triplets for triple signatures etc.). Finally, cells labelled as multiplets were isolated from the predicted scRNAseq data using the FilterMultiplet data with the default parameters (minCells = 2, minFreq = 10) and were utilized for a cell-cell interaction network plot using the PlotNetwork function with the default parameters (network_df, node_size = 20, node_color = ‘blue’, node_text_size = 4, edge_width_factor = 50, edge_color = ‘red’, network_layout = ‘fr’, legend_title = ‘scaled counts’, legend_position = ‘bottom’, min_edge_width = 0.5, max_edge_width = 3, main = ‘Network plot’, main_size = 15, hjust = 0.5, legend_text_size = 12, legend_title_size = 14).

Predicted doublet cells of T cell-Macrophage (T-Mac) interactions together with their singlet counterparts (single-positive T cells and Macrophage), were isolated and utilized for downstream analysis.

### 2.3 Isolation of T-Macrophage spatial spots

To identify T-Macrophage spatial spots (T-Mac spots) in the ovarian cancer spatial transcriptomics data, each spatial spot was tested for the colocalized enrichment of T cells and macrophages. Spots which simultaneously contained more than 10 % T cells and more than 10 % macrophages were considered colocalized T-Mac spots. T-Mac spots were identified for each clinical response group and normalised by the total number of spatial spots per group.

### 2.4 Functional signature enrichments

T cells, macrophages and T-Mac doublets were assessed for the enrichment of key T-cell and macrophage functionality programs. T-cell Functional signatures were obtained from the SignatuR package [29] and modified to include M1 macrophage, M2 macrophage, T-cell stemness, cell cycle, immune checkpoint, T-cell cytotoxicity, and T-cell exhaustion programs (supp. table 1). The UCell R package [30] was utilized to score cells for the functional signatures.

### 2.5 Differential expression of genes

Differential expression of genes (DEG) analysis was performed using the FindMarkers function from the Seurat package v5.1.1 in R v4.2.2, with the default parameters (slot = “data”, logfc.threshold = 0.1, test.use = “wilcox”, min.pct = 0.01, min.diff.pct = -Inf, only.pos = FALSE, max.cells.per.ident = Inf, random.seed = 1, latent.vars = NULL, min.cells.feature = 3, min.cells.group = 3). Genes were considered significantly differentially expressed if the adjusted p-value (p-adj) was less than 0.05. DEG was performed for doublet versus doublet or doublet versus singlet. For doublet versus singlet DEG, T cells and macrophages of the compared phenotype and response groups were merged to form a combined singlet population prior to comparison with the corresponding T-Mac doublets.

### 2.6 Pathway analysis

Gene set enrichment analyses (GSEA) were performed, using the log2FC values from DEG analysis, with the clusterProfiler R package. Enriched pathways were considered significant if the adjusted p-values < 0.05.

### 2.7 Transcription factor activity inference

Transcription factors (TF) activities were inferred using the decoupleR R package. This implements a univariate linear model to estimate whether a TF is active or inactive by evaluating the expression patterns of its target genes in the dataset of interest. The method leverages the CollecTRI (collection of transcriptional regulatory interactions) resource, a comprehensive collection of manually curated TF–target regulatory networks compiled from 12 independent databases and distributed through OmniPath. For each sample or cell and for every TF represented in the CollecTRI network, decoupleR fits a univariate linear model and generates a corresponding TF activity score. For computational efficiency, expression profiles were pseudobulked within each cell class (T-Mac doublets, macrophages, and T cells) and clinical groups using the aggregate.Matrix function of the Matrix.utils R package. TFs with p values < 0.05 were significantly enriched; the top 25 significant TFs for each pseudo sample cluster were taken and then visualised in a heatmap across all clusters. Furthermore, to obtain differential TF activities between cell clusters across singlet and doublet clinical groups, decoupleR was run against the log2FC values of significant DEGs (see section 2.3) from the compared groups. The top differentially active and inactive TFs were taken for each of the compared cell classes.

### 2.8 Ligand-receptor analysis

To assess ligand-receptor coexpression by T-Mac doublets, a list of ligand-receptor pairs (LRP) from Ramilowski et al. [31], containing 708 unique ligands and 691 unique receptors was obtained. For each T-Mac doublet, a ligand or receptor gene was considered expressed if its expression level in that doublet was higher than its mean expression across all T-Mac doublets. Hence, an LRP would be assumed to be co-expressed in a T-Mac doublet if both ligand and receptor genes were expressed in that doublet. The LRPs which were co-expressed in at least 5 doublets were retained for each clinical response group. The total number and proportion of doublets expressing each LRP was computed. Next, the co-expressed LRPs in the T-Mac doublets were validated for possible spatial colocalization in T-Mac spots (see section 2.3). For each spatial T-Mac spot, a ligand or receptor gene was considered expressed if the expression level was higher than the average expression across all spatial T-Mac spots. Similarly, an LRP is presumably colocalized in a T-Mac spot if both ligand and receptor genes are expressed in that spot. The number and proportion of spots expressing each doublet-enriched LRP was computed. Finally, a correlation analysis was performed to determine the relationship between the expression patterns of LRPs in T-Mac doublets and those of T-Mac spots.

## 3.0 RESULTS

### 3.1 scRNAseq reveals cellular landscapes in ovarian cancer patients

We obtained a fully processed and quality controlled scRNAseq dataset, with doublets removed, from Zheng et al. [24]. This dataset comprises ∼220,000 single cells from 13 treatment-naive ovarian cancer patients who underwent surgery followed by platinum-based chemotherapy. Patients whose disease progressed within 6 months after the completion of adjuvant therapy were classified as “resistant” while the others were classified as “sensitive”. Cell annotations revealed 10 distinct cell type clusters (Figure 1a), including B cells, CD4+ T cells, CD8+ T cells, NK cells, monocytes, macrophages, dendritic cells (DC), endothelial cells, cancer (epithelial) cells, and stromal cells. Group-level comparisons of cell-type composition showed a higher proportion of epithelial (cancer) cells in the sensitive group (∼27%) compared with the resistant group (∼14%) (Figure 1b & Supp. Table 1).

**Figure 1:**
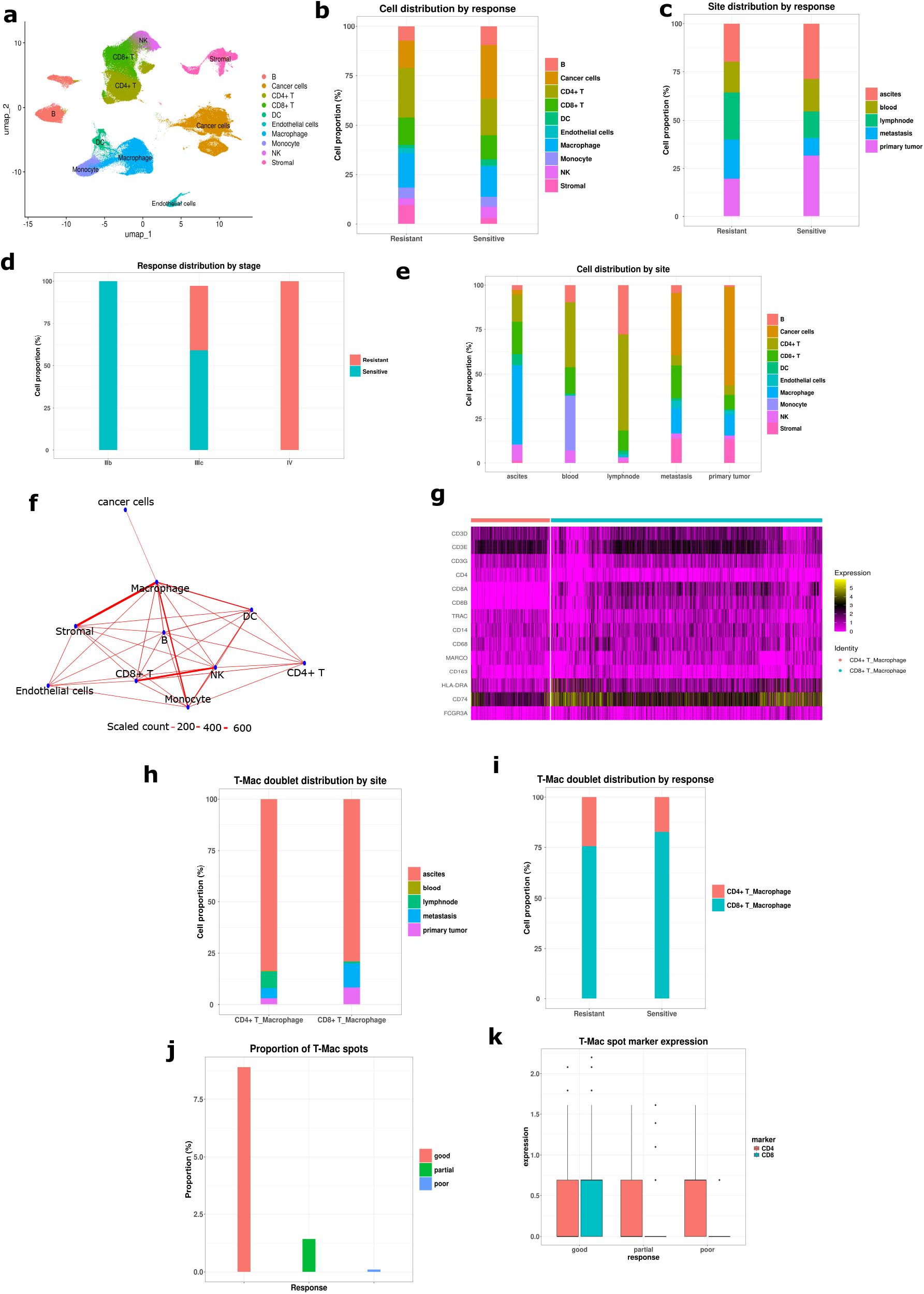
single-cell profiling of the ovarian cancer microenvironment, cell-cell interaction networks, and T cell-Macrophage doublet identification. A fully preprocessed scRNAseq data from 14 ovarian cancer patients with doublet excluded, as well as the corresponding unprocessed count matrices with doublets present, were obtained and profiled. ULMnet was used to infer physical cell-cell interaction networks from the unprocessed data. T cell-macrophage (T-Mac) doublets were identified and isolated from the inferred network. a) Annotated single-cell UMAP clusters showing 10 distinct cell types from the fully preprocessed scRNAseq dataset. b) Cell distribution in different therapeutic response groups. Resistant patients have lower cancer cell population but more macrophages and stromal cells. Sensitive patients have more B cells and NK cells. c-d) Cell distributions by sample sites and stages. Resistant patients have more metastatic and advanced stage cancers. e) Cell distributions by sample sites. Primary and metastatic tumor sites are dominated by cancer cells. Lymph nodes, ascites blood are populated by immune cells. Monocytes are exclusive to the blood. f) ULMnet generated physical cell-cell interaction networks from the unprocessed scRNAseq data (doublet present), using the annotated cell types in 1a for signature generation. g) A heatmap showing that T-Mac doublets co-expressed both T-cell and macrophage cell type-specific signatures h-i) T-Mac doublet subtype distributions by sample sites and therapeutic response. While both T-Mac subtypes dominate the ascites, CD8+ T-Mac doublets are more present at the primary and metastatic tumour sites. Also, sensitive patients have more proportion of CD8+ T-Mac doublets. j-k) Proportions and subtypes of spatial T-Mac spots identified from ovarian cancer spatial transcriptomics data. T-Mac spots representing spatially colocalised T cells and macrophages for potentially ongoing interactions, were generally more enriched in good responders, with the CD8+ subtypes being more important.

Clinical characteristics differed between response groups. Relative to sensitive cases, resistant cases included a higher proportion of metastatic samples, whereas sensitive cases included more primary tumour samples. (Figure 1c). Stage distributions also differed between groups: stage IIB cases were observed only in the sensitive group and stage IV cases only in the resistant group, while both groups included stage IIIC cases (Figure 1d). These differences were considered when interpreting group-level cellular composition. Hence, these show that even though the resistant patients presented with lower cancer cells, they possibly had more aggressive cancer cells that were potentially more difficult to treat. Across immune and stromal compartments, the resistant group showed higher proportions of CD4+ T cells, macrophages, and stromal cells, whereas the sensitive group showed higher proportions of B cells and NK cells. CD8+ T cell proportions were broadly similar between groups (Figure 1a; Supplementary Table 1).

We next examined cell-type composition across anatomical sites (Figure 1e). Primary and metastatic tumour sites were dominated by cancer cells, followed by stromal cells, with moderate immune infiltration. Ascites samples were enriched for macrophages, followed by CD4+ and CD8+ T cells, with a low fraction of cancer cells. Peripheral blood samples comprised immune populations, dominated by monocytes and CD4+ T cells. Lymph node samples were predominantly immune, with higher CD4+ T cell proportions, followed by B cells and CD8+ T cells.

### 3.2 Physical cell-cell interaction network

Using ULMnet [Ref], we inferred physical cell-cell interaction networks from the raw Zheng et al. scRNA-seq dataset in which doublets were retained [Ref]. After quality control and preprocessing (Methods), the doublet-retained dataset comprised ∼520,000 cells. Cell type-specific signatures were derived from the annotated singlet reference (Figure 1a) and applied to the doublet-retained dataset to identify multiplets and construct a physical interaction network (Figure 1f) . ULMnet generated a highly connected physical cell-cell interaction network with multiple immune-immune and immune-stromal cell interactions, with macrophages forming a highly connected node (Figure 1f).

Given the central role of macrophage-T cell coordination in antigen presentation and anti-tumour immunity, we focused subsequent analyses on predicted T cell-macrophage (T-Mac) doublets and their corresponding singlet populations (CD4+ T cells, CD8+ T cells, and macrophages).To assess whether predicted T-Mac doublets exhibited the expected composite transcriptional profiles, we examined canonical marker expression. Predicted T-Mac doublets showed concurrent expression of T cell markers (CD3D, CD3E, CD3G, TRAC) and macrophage markers (CD14, CD68, MARCO, CD163, CD74, HLA-DRA) (Figure 1g). (figure 1g). In addition, predicted CD4+ T-Mac doublets expressed CD4, whereas predicted CD8+ T-Mac doublets expressed CD8A and CD8B (Figure 1g).

In total, we identified 1323 CD8+ T-Mac and 384 CD4+ T-Mac doublets, predominantly from ascites. While CD8+ T-Mac doublets were reasonably more present in primary tumour and metastatic sites, the CD4+ T-Mac doublets were more largely found in the lymph nodes, with smaller fractions at tumour sites (Figure 1h). This underscores the importance of CD8+ T-Macs in the tumour microenvironment where cytotoxic cells are licensed to fight cancer cells. We next compared the relative composition of CD4+ versus CD8+ T-Mac doublets between clinical response groups. In both resistant and sensitive patients, CD8+ T-Mac doublets constituted the majority of T-Mac doublets. However, the resistant group displayed a higher relative fraction of CD4+ T-Mac doublets compared with the sensitive group (Figure 1i).

To validate the observed importance of T-Mac interactions in the tumour microenvironment (TME), we assessed T-Mac interaction in a spatial transcriptomics dataset of 8 ovarian cancer patients from Desienko et al. [26]. Patients were classified into 3 clinical groups as good, partial and poor responders. We first identified spatial spots where T cells were colocalised with macrophages (T-Mac spots), representing potential hotspots of ongoing T cell-macrophage interactions in the ovarian TME. We then profiled the spatial T-Mac spots in the context of therapeutic response, revealing that T-Mac spot populations correlated with response. Good responders displayed a remarkably high proportion of T-Mac spots which was about 5 times higher than seen in partial responders, while poor responders presented a negligible population of T-Mac spots (figure 1j). Finally, we assessed CD4- and CD8-associated gene expression within T-Mac spots across response groups. CD8+ T-Mac spots were exclusively found in good responders, presenting a balanced expression of CD4 and CD8. Partial and poor responders lacked CD8+ T-Mac spots, exclusively presenting only CD4+ T-Mac spots (Figure 1k).

Taken together, these results show that T-Mac interactions are relevant in the ovarian TME, with CD8+ T cells being more crucial to this interaction than CD4+ T cells for favourable response to chemotherapy.

### 3.3 Functional signature enrichments

To characterise functional profiles within predicted T cell-macrophage doublets (T-Mac), we scored T-Mac doublets using UCell across curated T-cell and macrophage gene signatures, including T-cell stemness, cytotoxicity, immune checkpoint signalling, exhaustion, cell-cycle program, and macrophage polarization (M1 and M2) signatures. Across all T-Mac doublets, enrichment scores differed between clinical response groups (Figure 2a). T-Macs from the resistant patients displayed more enrichment of T-cell stemness, cytotoxicity and cell cycle signatures than the sensitive group. However, they were concurrently enriched in T-cell exhaustion and the protumoral M2 macrophage signatures, lacking the antitumoral M1 macrophage signatures. On the other hand, the T-Mac doublets from the sensitive patients are enriched in the antitumoral M1 macrophage signature and relatively lack T cell exhaustion signatures, with reasonable levels of cytotoxic and proliferative activities (figure 2a). Although there was also M2 macrophage signature enrichment in sensitive T-Macs, the concomitant enrichment of M1 signatures and the absence of exhaustion signatures might have overwhelmed the influence of the M2 macrophages.

**Figure 2:**
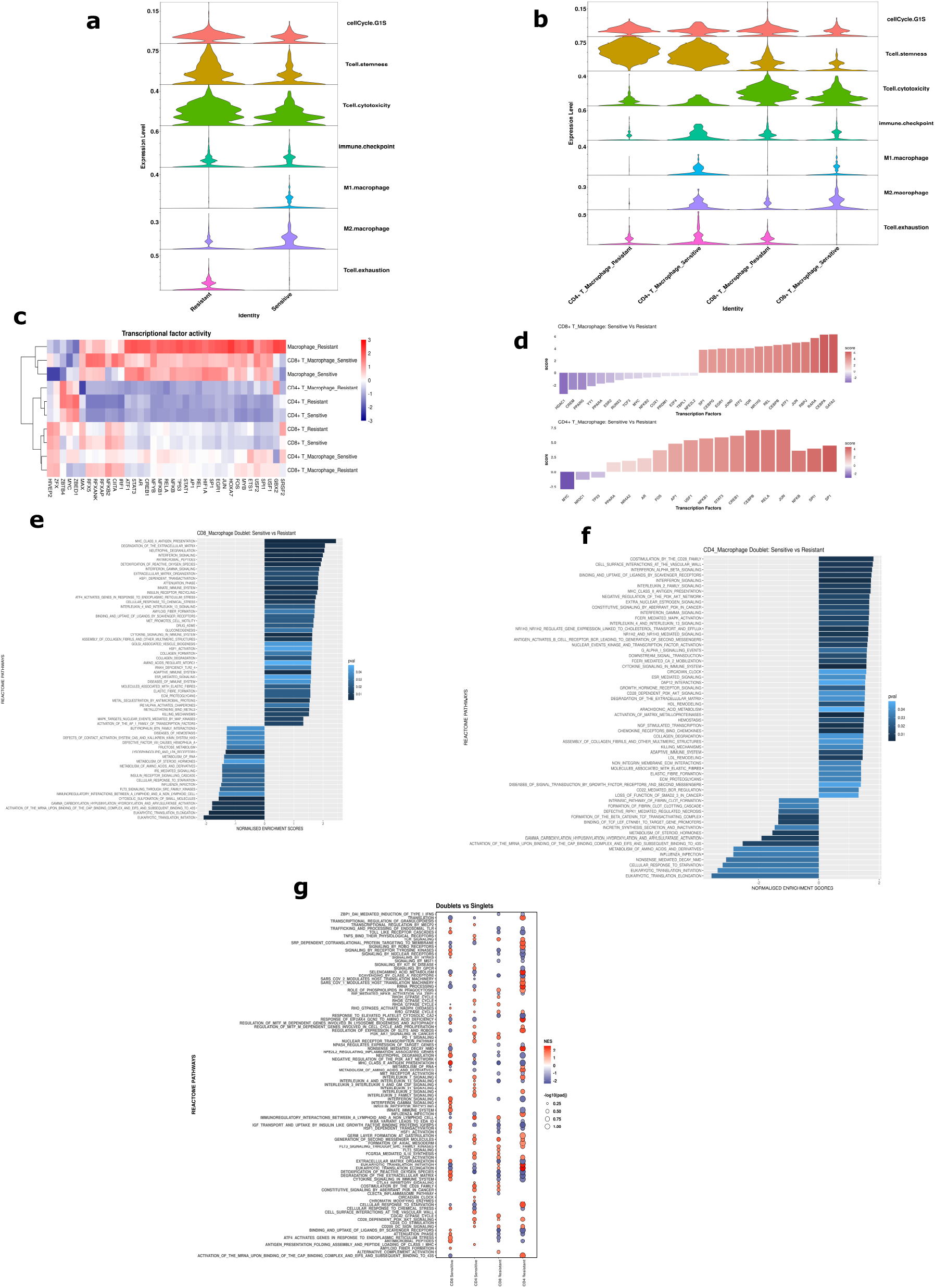
Signature enrichments, transcriptional factor inferences and functional annotation. a-b) Functional signature enrichments in T-Mac doublets. T-Macs in sensitive patients possess M1 macrophage signature and relatively lack T-cell exhaustion signature. T-Macs from resistant patients lack the M1 macrophage signature but possess the T-cell exhaustion signature. T-cell cytotoxicity signature was more enriched in CD8+ T-Macs as expected c) Transcriptional factor (TF) enrichment in singlet and T-Mac doublet populations from different therapeutic response groups. T-Mac doublets are widely separated by therapeutic response while singlets cluster by phenotype and therapeutic response. d) Differential TF activities between T-Mac doublets from sensitive and resistant patients for CD8+ T-Macs (top) and CD4+ T-Macs (bottom). Differential TF activities were inferred from the significantly differentially expressed genes (DEGs; p <0.05) between the compared groups. TFs were considered differentially active if p value was less than 0.05, the top 15 differentially active and inactive TFs were displayed. e-f) Functional annotation of DEGs when comparing T-Mac doublets from sensitive and resistant patients using the reactome database, for CD8+ T-macs (e) and CD4+ T-Macs (f). g) Functional annotation of DEGs when comparing T-Mac doublets to singlets (T cells and macrophages) from the same response group, using the reactome database.

We next stratified T-Mac doublets by T-cell phenotype (CD4+ versus CD8+) to determine which populations contributed to the observed enrichment patterns (Figure 2b). Stemness-associated scores were higher in CD4+ T-Mac doublets compared with CD8+ T-Mac doublets, whereas cytotoxicity-associated scores were higher in CD8+ T-Mac doublets (Figure 2b). M1 macrophage signature enrichment was observed in CD4+ and CD8+ T-Mac doublets from sensitive patients but was minimal in the corresponding resistant groups (Figure 2b). Notably, CD8+ T-Mac doublets from sensitive patients showed low exhaustion-associated scores relative to other T-Mac subsets (Figure 2b)

Altogether, these results emphasised the role of M1 macrophage and CD8+ T-cell interactions in driving favourable response in ovarian cancer patients.

### 3.4 T-Macrophage doublets display transcriptionally and functionally different states

A global analysis of TF activities on pseudobulk profiles revealed that T-Mac doublets segregated by therapeutic response, with CD8+ T-Mac and CD4+ T-Mac doublets showing distinct TF activity patterns between sensitive and resistant groups (Figure 2c). In contrast, TF activity profiles of singlet CD4+ T cells, singlet CD8+ T cells, and singlet macrophages were more similar across response groups than the corresponding doublets (Figure 2c). In addition, TF activity patterns differed between singlet T cells and singlet macrophages, consistent with their distinct lineages (Figure 2c).

We next performed differential TF activity analysis between sensitive and resistant T-Mac doublets (Methods). In both CD8+ and CD4+ T-Mac doublets, multiple TFs commonly associated with immune activation and signalling (including JUN/JUND, REL/RELA, NFKB1, EGR1, SP1, FOS, and ATF-family members) showed higher inferred activity in sensitive relative to resistant doublets (Figure 2d). This indicated that the interacting T cells in the sensitive group are more active and poised to perform effector T-cell functions. To explore possible doublet-specific TFs, we compared T-Mac doublets of each response group to their singleton T cells and macrophage counterparts. In general, T-Macs from resistant groups presented inactive TFs that included T-cell activation/functionality related TFs, compared to the singlets (supp. figure 1a-b). This possibly inferred that both CD4+ and CD8+ T cells from resistant patients underwent reduced activity following their interactions with macrophages due to immunosuppressive interaction events.

To identify pathways distinguishing T-Mac doublet states, we performed pathway enrichment analysis of DEGs between response groups (Methods). In CD8+ T-Mac doublets, sensitive versus resistant comparisons showed enrichment of interferon signalling (IFN-γ and IFN-α/β), cytokine signalling, and AP-1-related programmes among the top pathways (Figure 2e).

We next investigated the functional pathways that define T-Macs by performing functional annotations of the DEGs. Both CD8+ and CD4+ T-Macs from the sensitive patients showed differential enrichment of key immune related pathways compared to the CD8+ T-Macs from the resistant patients. This included interferon signalling, cytokine signaling, MAP kinases signaling, IL-2 signalling, interferon gamma signalling, CD28 signalling, and AP1 activation signaling pathways (Figures 2e & 2f). This further depicts the activation state of T-Macs from sensitive patients. We also compared T-Mac doublets of different phenotype (CD4+/CD8+) and response groups (sensitive/resistant) to their singlet counterparts. Compared to the T-cell and macrophage singlets, T-Mac doublets generally showed differential enrichment of TCR signaling, CD28 co-stimulation, IL-2 signaling, FCGR signaling, and phagocytosis (Figure 2g). This is a possible indication of the known series of events involved in antigen presentation leading to direct physical interaction between T cells and macrophages through the formation of immune synapses [32]. Briefly, macrophages ingest tumour antigens via phagocytosis, process the antigen and present to T cells in an MHC-TCR dependent manner, thereby inducing TCR signaling and downstream events that induces IL-2 secretion and T cell activation/proliferation. Upon deeper investigation, we observed that CD4+ and CD8+ T-Mac doublets from the resistant patients had downregulations of cytokines and interferon mediated pathways (Figure 2g). This further reinforces the observed immunosuppressive modulation of T-Macs from the resistant group.

Overall, these results depict that T-Macs were in distinct transcriptional and functional states that influenced patient response to chemotherapy.

### 3.5 Ligand receptor interactions depict ongoing T-Macrophage interactions

We carried out ligand-receptor interactions by assessing co-expressed ligand-receptor pairs (LRP), using a list of ligands and cognate receptors obtained from Ramilowski et al. [31]. Briefly, a ligand or receptor gene is considered expressed in a given T-Mac doublet if the expression level of that gene is greater than its average expression across all T-Mac doublets. We then consider an LRP to be enriched if it is present in at least 5 T-Mac doublets, identifying 280 enriched LRPs from sensitive T-Macs and 398 enriched LRPs from resistant T-Macs (supp. table 1). Of these, 273 LRPs were commonly expressed by both sensitive and resistant T-Macs (supp. table 1). We found out that T-Macs generally co-expressed LRPs that potentially mediate ongoing antigen presentation processes. This included key human leukocyte antigen (HLA) and T-cell receptor (TCR) interaction pairs such as HLA-A_CD3G, HLA-C_CD3G, HLA-B_CD3D, HLA-B_CD3G, HLA-A_CD3D, B2M_CD3D and B2M_CD3G (figures 3a & 3b, supp. table 1). These LRPs are indicative of ongoing antigen presentation where macrophages present peptide antigens in the context of HLA molecules to T cells by physical contacts via TCR molecules on T-cell surfaces . Furthermore, there was enrichment of several LRPs involving integrins that are classically associated with immune synapse formation to stabilize cell-cell contacts and facilitate TCR signalling. These LRPs included several LFA-1 (αLβ2) and VFA-4 (α4β1) family integrins and their respective ICAM and VCAM ligand family proteins [33] such as ICAM1_ITGB2, ICAM1_ITGAL, ICAM2_ITGB2, ICAM2_ITGAL, ICAM3_ITGB2, ICAM3_ITGAL, VCAM1_ITGA4, VCAM1_ITGB1, VCAM1_ITGB7 (figures 3a & 3b, supp. table 1). These integrins are also essential to form and constitute the peripheral supramolecular activation clusters (p-SMAC) that stabilize T-cell-APC contact, sustain TCR signalling, provide co-stimulatory signals, and promote T-cell activation, differentiation and effector function [34]. We validated these findings in the spatial transcriptomics data by evaluating spatial enrichment of the identified LRPs (from T-Mac doublets) in spatial T-Mac spots. Again, a tested LRP was considered expressed in a T-Mac spot if both ligand and receptor genes are expressed at a level higher than the average expression across all T-Mac spots (see methods). Indeed, 253 of the 280 LRPs enriched in T-Mac doublets from sensitive patients were enriched in at least 5 T-Mac spots from good responders (supp. table 1). Also, 209 of the 398 LRPs from resistant T-Macs were enriched in at least 5 T-Mac spots from bad (partial and poor) responders (supp. table 1). Interestingly, the enriched LRPs in the T-Mac spatial spots generally included LRPs that are relevant for antigen presentation and immune synapse formation as seen in T-Mac doublets above. Furthermore, we found a reasonably positive correlation between the expression patterns of enriched LRPs in T-Mac doublets and spatial T-Mac spots (figures 3c & 3d) for both sensitive patients/good responders (r = 0.33) and resistant patients/bad responders (r = 0.27). These results reconfirm that T-Mac doublets represent active physical cell-cell interaction units of ongoing antigen presentation and T-cell modulation.

**Figure 3:**
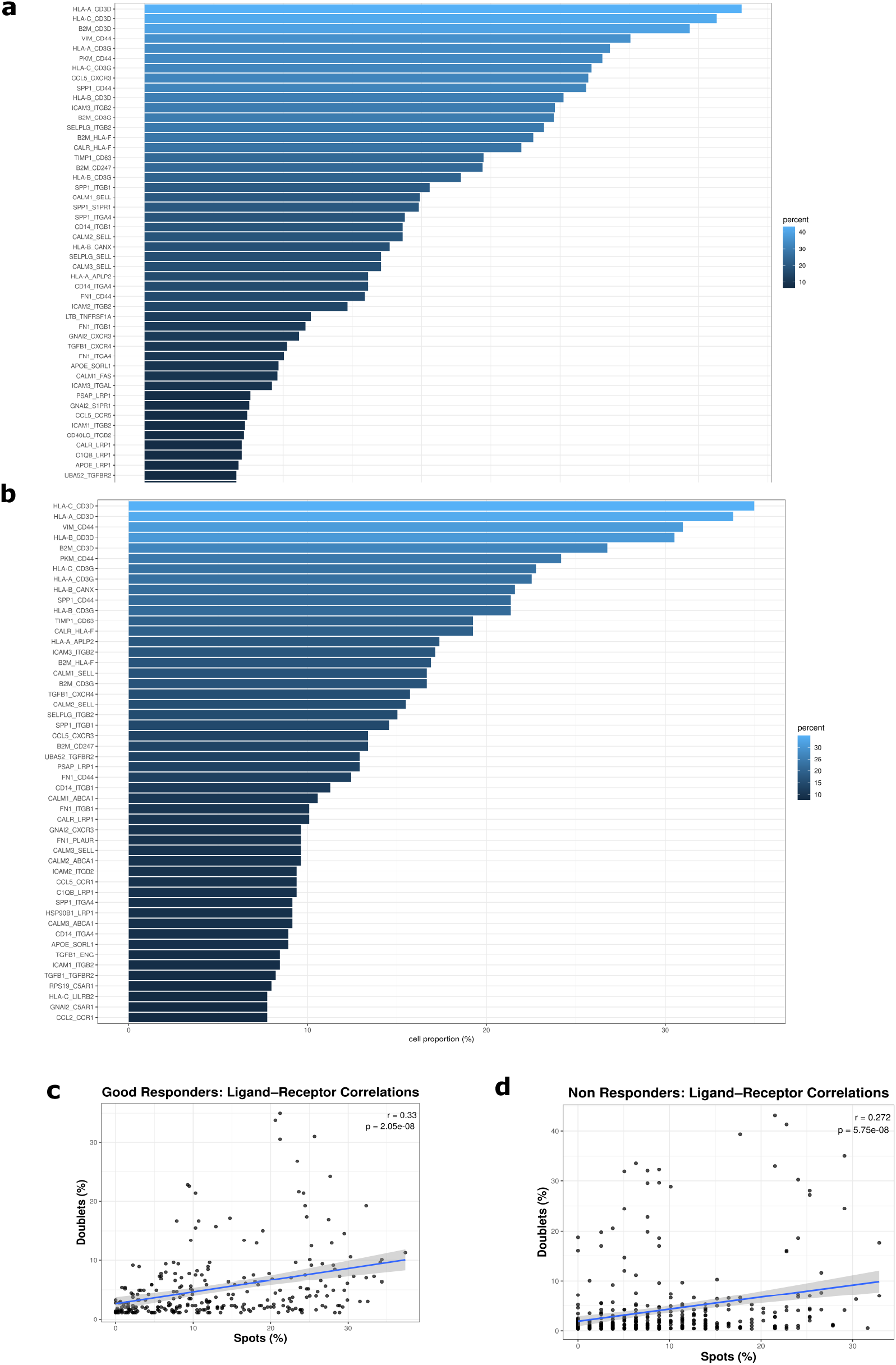
Ligand-receptor analyses. a-b) Enriched ligand-receptor pairs (LRP) in T-Mac doublets from (a) sensitive and (b) resistant patients. A ligand or receptor gene is considered expressed in a T-Mac doublet if the expression level in that doublet is greater than the average expression across all doublets. An LRP is considered enriched in a doublet if both ligand and receptor in that pair are expressed in at least 5 doublets. Several LRPs involving cell-cell contact for antigen presentation were generally enriched in T-Mac doublets. c-d) Correlations between LRP enrichments in T-Mac doublets and spatial T-Mac spots. A ligand or receptor gene is considered expressed in a spatial T-Mac spot if the expression level in that spot is greater than the average expression across all T-Mac spots. An LRP is considered enriched in a doublet if both ligand and receptor in that pair are expressed in at least 5 T-Mac spots. Doublet-level and spot-level enrichments were compared using Pearson’s correlation.

## 4.0 DISCUSSION

Doublets in scRNAseq are recently being considered a good source of biological information to study physical cell-cell interactions in tissues [20] [21]. In this study, we applied ULMnet to an ovarian cancer scRNA-seq cohort and focused on macrophage-T cell interactions to explore their association with therapeutic response, with additional evaluation in an independent spatial transcriptomics cohort..

Initial profiling indicated that clinical and sampling characteristics differed between response groups, including differences in stage and the distribution of primary versus metastatic samples. These factors likely influence tumour composition and immune infiltration and therefore represent important context when interpreting response-associated differences in cell-type proportions. Notably, the resistant patient group showed a higher proportion of stromal cells, consistent with prior evidence that stromal programmes, particularly fibroblast-associated signalling, can support tumour progression, metastatic potential, and immunomodulation within the tumour microenvironment [35] [36]. In addition, the sensitive patient group showed higher NK-cell representation, consistent with established roles for NK cells in anti-tumour immunity through cytotoxic effector mechanisms and cytokine production [37].

Applying ULMnet to the doublet-retained dataset produced a highly connected interaction network reflecting extensive immune-immune and immune-stromal contact patterns in ovarian cancer.. Focusing on T-Mac doublets, we observed response-associated differences in the relative abundance of CD8+ versus CD4+ T-Mac doublets, with CD8+ T-Macs relatively enriched in sensitive patients and CD4+ T-Macs relatively enriched in resistant patients. Importantly, these trends were supported in spatial transcriptomics, where spatial CD8+ T-Mac spots were found to be remarkably higher in good responders. This indicates that CD8+ T cells are frequently colocalised with macrophages and are potentially engaging in physical interactions with the macrophages in good responders. CD8+ T cells are the backbone of anticancer immunity where they directly target and kill cancer cells expressing cancer antigens after being primed by APCs such as macrophages [38]. Therefore, the CD8+ T cells engaging macrophages in CD8+ T-Macs would eventually target and kill cancer cells after such interactions, thereby contributing to positive therapeutic outcomes. Previous studies have shown that higher CD8+ T cell proportion in ovarian cancer contribute to better survival outcomes [39] [40] [41] [42], hence the higher enrichment of CD8+ T-Mac in sensitive patients, and higher population of CD8+ spatial T-Mac spots in good responders, promote more tumour killing, thereby enhancing positive therapeutic response. Apart from T-cell phenotypes in T-Macs, we identified that macrophage phenotypes also impacted therapeutic outcomes. T-Macs in sensitive patients involved the engagement of antitumoral M1 macrophage phenotypes whereas the macrophages in the resistant T-Macs were predominantly of the protumoral M2 phenotypes. M1 macrophages are classically activated macrophages which possess antitumoral properties including inflammatory response, phagocytosis of tumour antigens, and antigen presentation to T cells for effector tumour killing. They also secrete immunomodulatory mediators that promote T-cell activation and strengthen T-cell function [43]. This infers that the T cells in these interactions are potentiated for tumour killing, thereby contributing to the observed favourable response to chemotherapy in the sensitive patients. On the other hand, M2 macrophages are alternatively activated macrophages that possess protumoral properties, secreting mediators that induce immunosuppression, thereby dampening T-cell mediated response and promoting tumour progression [43]. Therefore, despite the cytotoxic, proliferative and self-renewing properties of the T cells in resistant patients, they are engaged with M2 macrophages which modulate their functions thereby inducing exhaustion. This reduced potential tumour killing, promoting tumour progression and could be a contributing factor to the observed aggressiveness of the cancer cells in these patients and therapeutic resistance. Indeed, a higher proportion of M1 polarized macrophages have been reported to favour positive therapeutic outcomes in high grade serous ovarian cancer patients, with longer overall survival and progression free survival [44]. Since M1 macrophages act by secreting proinflammatory cytokines and antitumoral mediators, their interactions with T cells in T-Macs would induce T-cell activation and effector function. This effect was observed at transcriptional factor level where the differentially active TFs in T-Macs from sensitive patients compared to resistant patients were those involved in T-cell activation, proliferation and effector function. On the other hand, M2 polarized macrophages are known to favour chemotherapeutic resistance in ovarian cancer through the secretion of protumoral factors which reduce drug accumulation at tumour sites, induce resistant gene expression, and provide prosurvival signals [45]. In addition, M2 macrophages secrete immunosuppressive mediators that dampen immune responses, leading to T-cell exhaustion and metastatic proliferation [43]. Therefore, these dual roles of M2 polarized macrophages on cancer cells and T cells via immunosuppressive T-Mac interactions have negatively contributed to the therapeutic resistance in the resistant patients.

Since macrophages are professional APCs and are only expected to contact T cells physically for the sole purpose of antigen presentation, we demonstrated that T-Mac doublets were specifically enriched in antigen presentation and co-stimulation related pathways. This was reinforced by ligand-receptor interactions involving HLA/TCR pairs including several LFA-1/ICAMs and VFA-4/VCAM integrin interactions which demonstrated ongoing antigen presentation and immune synapse formation.

Although our analysis revealed clinically significant results, we do have some key limitations. Firstly, we inferred physical cell-cell interactions from scRNAseq doublets with the assumption that all predicted doublets are true interaction events arising from unseparated physically connected cells. However, some random co-encapsulation of non-interacting cells often occurs in scRNAseq, generating random doublets. Therefore, it is possible that some of the characterized doublets might be random events rather than true biological doublets of unseparated physically connected cells. Nonetheless, ULMnet assigns enrichment scores to predicted doublets and the associated threshold might have partly eliminated the random doublets. Also, the correlation between our doublet models and spatially co-localised cell pairs, yielding biologically and clinically plausible results, further demonstrate that the analysed doublets are not just spurious interactions but possess high likelihood of being true T-Mac doublets. Secondly, we had a low starting number of T-Mac doublets for each therapeutic response group, due to the stringency of the scRNAseq protocol, and we analysed doublets derived only from 13 patients, this limits the generalizability of our conclusions. Future research to include more patients and a higher number of T-Mac doublets would solidify the robustness of our findings. Finally, we presented results arising from in silico analysis of physical interactions. Although this was partly strengthened by spatial transcriptomics dataset, experimental validations are required to consolidate our findings. For example, T-Mac doublets derived from in vitro co-cultures or in vivo derived pairs from partially dissociated and FACS sorted samples could be complementarily analysed. Also, key genes and LRPs inferred from our analysis can be validated from orthogonal approaches such as qPCR, and fluorescence in situ hybridization.

In conclusion, we have demonstrated that T cell-macrophage interactions in the ovarian cancer microenvironment influence therapeutic outcome in ovarian cancer patients. These outcomes were dependent on T-cell and macrophage phenotypes, with CD8+ T cells and M1 polarized macrophage interactions being more favourable for positive therapeutic outcomes.

## 5.0 ACKNOWLEDGEMENT

Financial support was received for this work and/or its publication. Science Foundation Ireland through the SFI Centre for Research Training in Genomics Data Science under 18/CRT/6214, Strategic Partnership Programme Precision Oncology Ireland co-funded by SFI and AstraZeneca (18/SPP/3522), and Health Research Board, Ireland (ERATRANSCAN-2022-001, EPPerMed-2024-1).

## 6.0 DATA AVAILABILITY

The publicly available scRNAseq and spatial transcriptomics datasets with the associated metadata used for this analysis are available in the cited studies.

All scripts and intermediate files associated with this analysis will be made available upon reasonable request through the corresponding author.

---

**Supplementary figure 1.**

a) Differential TF activities between CD8+ T-Mac doublets and the corresponding singlets from sensitive (top) and resistant (bottom) patients. Differential TF activities were inferred from the significantly differentially expressed genes (DEGs; p <0.05) between the compared groups. TFs were considered differentially active if p value was less than 0.05, the top 15 differentially active and inactive TFs were displayed.

b) Differential TF activities between CD4+ T-Mac doublets and the corresponding singlets from sensitive (top) and resistant (bottom) patients. Differential TF activities were inferred from the significantly differentially expressed genes (DEGs; p <0.05) between the compared groups. TFs were considered differentially active if p value was less than 0.05, the top 15 differentially active and inactive TFs were displayed.

## Notes

### Competing Interest Statement

The authors have declared no competing interest.

