## Supplementary figures and images for "T cell-Macrophage Interactions Potentially Influence Chemotherapeutic Response in Ovarian Cancer Patients"

### Supplementary Figure 1

#### Supplementary figure 1

a

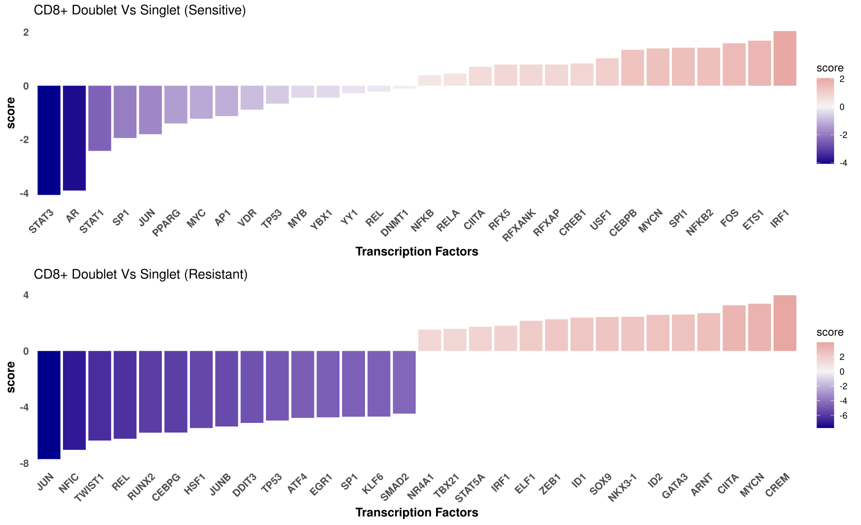


b

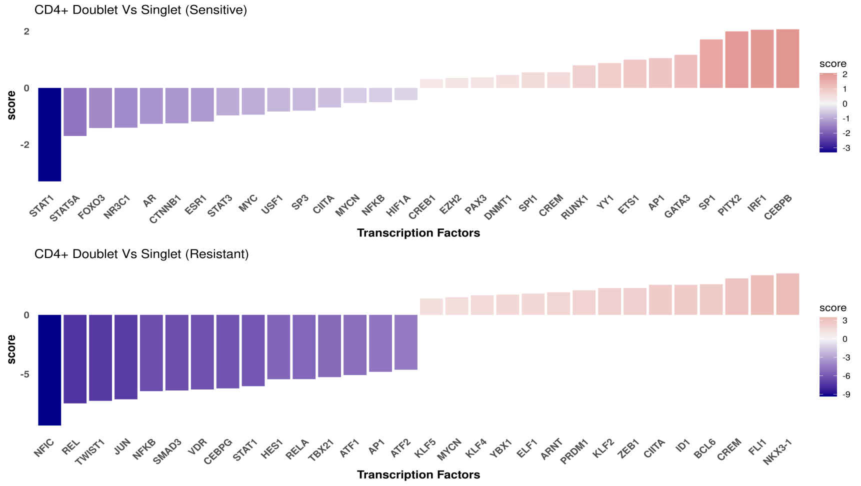
